# Neighborhood size-effects in nonlinear public goods games

**DOI:** 10.1101/347401

**Authors:** Gregory J. Kimmel, Philip Gerlee, Joel S. Brown, Philipp M. Altrock

## Abstract

Ecological and evolutionary dynamics can be strongly affected by population assortment and frequency-dependent selection. In growing populations, a particular challenge is to disentangle global ecological effects from local frequency-dependent effects. Here we implement a logistic growth and death model on the global scale, coupled to frequency-dependent growth rates influenced by a public goods game between cooperators and defectors. For each individual, the public good is only effective within a neighborhood of other individuals, and the public good-growth rate relationship can be nonlinear. At low numbers of cooperators, increases of public good accumulate synergistically; at high numbers, increases in public good only provide diminishing returns-the inflection point of this pattern is given by the strength of frequency-dependent selection in relation to the background fitness effect. We observed complex critical behavior in the evolutionary dynamics’ equilibria, determined by the relative magnitude of frequency-dependent to constant (background) growth benefits. We predict neighborhood-size-driven state changes, hysteresis between polymorphic and monomorphic equilibria, and observed that type-dependent differences in neighborhood sizes can destabilize monomorphic cooperative states but increase coexistence of cooperators and defectors. Stochastic neighborhood size fluctuations also led to coexistence and could stabilize the purely cooperative equilibrium. Our results quantify the role of assortment through neighborhood-size effects and nonlinearity of the gains function in eco-evolutionary dynamics, which is relevant for a variety of microbial and cellular public goods games.

---

Nature offers diverse examples of individuals providing public goods that benefit the provider as well as others nearby. Numerous mammal species provide alarm calls or group vigilance against predators. In banded mongooses, an individual in the foraging line may scare up insects for others [1]. Prairie dogs clear brush and open up sight lines around their burrows that become available to others in the colony [2]. Predator inspection by guppies [3], or mobbing of hawks by crows [4] provide useful information and safety for others, regardless of whether one participates in the inspection/mobbing or not. Colonial spiders contribute to ever larger and successful webs. Via an associational refuge, a plant can defend against herbivores using spines or toxins, which can provide protection for itself and nearby unprotected plants by dissuading the herbivore from feeding in some neighborhood of the unpalatable plant [5, 6]. Microbes provide public goods either by secreting defensive chemicals, as in the case of bacterial biofilms or by co-feeding [7, 8]. Yeast and other protists synthesize and spill essential nutrients, such as key amino acids into their surroundings [9]. These examples highlight the range of potential mutually beneficial interactions across multiple scales.

Cancer cells, too, behave as communities [10] and engage in public goods games [11, 12, 13]. Within a tumor, cells compete for resources, but also act as ecological engineers [14]. Through individual and collective actions, tumor cells can engineer a more favorable environment [10], which has led to the suggestion that cancer cells may evolve cooperation [15, 16], and that normal cells and cancer cells engage in a mutualistic relationship [17]. It may be strictly beneficial to a cancer cell to recruit blood vessels, signal normal cells and defend against the immune system, but these actions likely benefit neighbors too. As soon as there are shared benefits, but an individual’s cost to the production of the public good, an evolutionary game emerges. Such public goods games can strongly influence the eco-evolutionary dynamics between cooperators (or producers) and defectors (or consumers/free-riders).

All of the above cases represent social dilemmas based on the defector problem. They have been approached as variants of the snowdrift game [18] or other producer-scrounger games [19, 20]. Collective fitness or collective payoffs are maximized when everyone produces; yet often the evolutionarily stable strategy (ESS) of these games has either a mixture of cooperators and defectors; or no cooperators at all. Various models and hypotheses have been proposed to explain how evolution may favor a higher frequency of cooperators than might otherwise be expected. Three common hypotheses include 1) non-random assortative interactions where like interacts with like more frequently than by chance [22, 23], 2) iterative plays of the game where players can condition their actions on whether others cooperate or not, and 3) kin selection [21].

Here we model features of public goods games with the goal of expanding their utility and rendering them more applicable to populations within their natural environments. In particular, we see the game as embedded within the larger context of a population’s eco-evolutionary dynamics. Whether we examine prairie dogs, yeast [24], or cancer cells [25], the population exhibits ecological dynamics in form of changes in population size, and evolutionary dynamics in the form of changes in the frequency of producers relative to consumers. We see the former as density effects and the latter as frequency effects [26]. Here we integrate density and frequency dependence into a single modeling framework.

Public goods game formulations often assume a linear relationship between benefits and the number of cooperators [27, 28]. In many situations, however, the collective benefit may be non-linear with respect to the number of cooperators [29, 30, 31]. If one alarm call is sufficient to alert the colony of prairie dogs to a predator, then there are rapid diminishing returns to having several callers. If a collective defense by microbes against a predator, or by cancer cells to the immune system, requires a threshold effect, then there may be increasing returns where two producers more than double the collective benefit. We are particularly interested in the consequences of having a non-linear relationship between the collective benefit and the number of cooperators.

Benefits that are produced and shared are often only available within a finite neighborhood [32, 33]. These finite neighborhoods create a form of assortative interaction: an individual always experiences a slightly higher frequency of its own strategy within a neighborhood than present in the entire population. This tilt is because the individual contributes to the public good if it is a cooperator, and detracts from the collective public good if it is a defector. Consider a single individual in a neighborhood. The very fact that it is either a cooperator or a defector biases that individual’s experience with other cooperators and defectors. Self and non-self interactions can lead to assortment even when all neighborhoods are established randomly. In particular, cooperators may interact more locally than defectors. We are interested in how strategy-specific neighborhood sizes influence the population dynamics and the ESS, especially in terms of promoting polymorphic populations of cooperators and defectors which coexist.

## Nonlinear public goods in a growing population

In this section we present the basic features of our mathematical model. We consider the dynamics of a population that consists of two sub-populations. The model is general and applies to ecological systems at a number of different scales. However, in the following, we imagine cells that produce a growth factor which acts as a public good. Producer cells of type *C* (as in cooperators) produce a public good at a cost and consume it to gain a benefit. Consumer cells, type *D* (as in defectors), consume the public good, if available, but do not produce nor incur a cost. Each cell type’s intrinsic growth rate increases with the availability of the public good. We assume type-specific death (or apoptotic) rates *δ*_*C*,*D*_, and that cells obey a logistic growth law with an overall carrying capacity *K*. The densities of cooperators *C* and defectors *D* are given by *x*_*C*_ and *x*_*D*_. Changes in densities can be described by the dynamic system

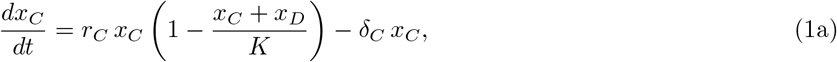

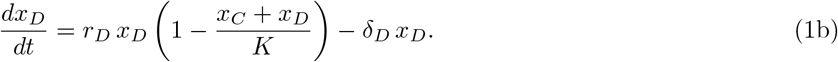

We are interested in cases where the intrinsic growth rates, *r_C_* and *r_D_*, depend on the frequency of cooperators *y* = *x*_*C*_/(*x*_*C*_ + *x*_*D*_).

We assume that the public good is a local commodity, for example a growth factor produced by *C*, and consumed by all within a certain neighborhood. We assume that growth factor production brings a direct benefit that amounts to an increased intrinsic growth rate, measured by the benefit parameter *β*. Each cell can benefit from the public good produced in a neighborhood of *n* cells. If the overall fraction of producer cells is *y*, and the population is well mixed, we can randomly assemble the neighborhood of *n* cells to obtain the expected public good-related benefit:

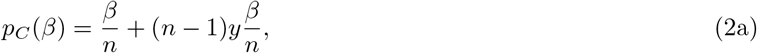

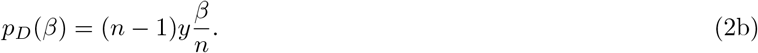

These expectation values result from binomial sampling *n* − 1-times with a success probability of *y*. A focal *D* cell can only be in a neighborhood of maximally *n* − 1 other cells (of which all could be cooperators). Cooperators generate a benefit-to-self. At this level, *C* always receives a higher benefit than *D*, but *C*’s bear a cost that directly reduces their intrinsic growth rate.

We assume that the benefit to each member of a neighborhood may increase non-linearly with the frequency of *C* within the neighborhood [13, 14, 29]. For example, one could observe steep increases of fitness at low frequencies of *C*, followed by a saturation effect in fitness once the fraction of *C* crosses a threshold. To describe such a complex nonlinear relationship mathematically we require at least two parameters. A sigmoidal, or s-shaped fitness curve can be described using the already introduced parameter *β*, and a frequency independent (background) benefit-parameter *σ*. The inflection point in which benefit increase is maximal is then given by *σ*/*β*. If the fraction of *C* is below *σ*/*β*, additional cooperators create synergies. Above *σ*/*β*, the returns from adding cooperators to the neighborhood diminishes. Thus, a sigmoidal-shaped curve describes the relationship between frequency of cooperators and the benefit to an individual. Including the cost for cooperation, *κ*, we write the frequency-dependent intrinsic growth rates in the following normalized form

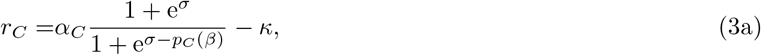

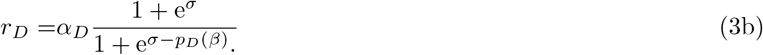

A sketch of the interaction between *C* and *D* via public good production is provided in Figure 1 A,B. Without any benefit from cooperators, *β* = 0, we obtain the intrinsic growth rates *r*_*C*_ = *α*_*C*_ − *κ* and *r*_*D*_ = *α*_*D*_. For a very small benefit, and allowing σ to be arbitrarily large (no upper limit on the positive effect of additional public good production), we can linearize the intrinsic growth rates *r*_*D*_ = *α*_*C*_(1 + *pC*(*β*)) - κ, and *r*_*D*_ = *α*_*D*_(1 + *pD*(*β*)). Linear relationships between benefits and the amount of the public good have frequently been used in recent models [25, 28, 34, 35].

In addition to neighborhood size *n*, we wanted to study the effects of strategy-specific neighborhood sizes. Cooperators may have a neighborhood size of *n*_*C*_, and defectors a neighborhood size of *n*_*D*_. These two different neighborhood size parameters could take any positive values. We specifically considered the possibility of neighborhood size differences that would make it possible for *C* to invade *D*, i.e. *n*_*C*_ < *n*_*D*_. Cooperators should favor smaller neighborhood sizes to increase their share of the public good that they contribute. Whereas defectors should favor larger neighborhood sizes as a way to dilute their failure to contribute to the public good.

We also wanted to explore fluctuations in *n*. Instead of considering *n* as an exogenous parameter, we defined a fixed number *N* of compartments among which all cells are randomly distributed to form neighborhoods. In this way neighborhood sizes would vary according to a multinomial distribution with a mean of (*x*_*C*_ + *x*_*D*_)/*N*. To simulate this, we ran a dynamic loop of two steps. First, we evaluated the outcome of a public goods game and logistic growth within each compartment for a fixed amount of time Δ*t*, according to (1) with *n* and *y* as compartment-specific quantities. Second, we modeled cell migration by pooling the individuals from all compartments and re-distributing them randomly among the compartments.

For all of the above cases, we examined how the nonlinear benefit-growth rate relationship and the neighborhood size(s) determine the number and stability properties of the equilibria (ESSs). We found cases of hysteresis, where the dynamics gives rise to alternate stable states, in the form of two or more ESSs that are locally convergent stable, depending upon the initial conditions. Given a fixed cost of public good production, the three key parameters that determine the number and stability of the eco-evolutionary system are the neighborhood size *n* (or *n*_*C*, *D*_), and the benefit parameters σ and *β*. All mathematical and computational tools and proofs are presented in the Methods section.

## Results and Discussion

How do independent and dependent effects of the nonlinear public good influence the dynamics of cooperators? For large neighborhood sizes, the public good essentially becomes global. This global frequency-dependence typically leads to the tragedy of the commons, in which cooperative producers cannot prevail. Frequency-dependent and-independent effects of the public good, in sufficiently small neighborhoods, can counteract the tragedy of the commons, because local accumulation benefits the cooperators. In the following we discuss key properties of our nonlinear public goods model, including how polymorphic equilibria can emerge and how they behave under changes to the benefit function. To this end, we also infer the possible shape of a frequency-dependent benefit function from a previously published cancer cell growth-experiment [13]. Lastly, we are interested in how strategy-dependent neighborhood sizes, and fluctuation in neighborhood sizes can strengthen coexistence of defectors and cooperators.

### Escaping the tragedy of the commons

How do growth rate differences allow cooperation to persist? If the cost of cooperation is exorbitantly high, the all-*D* state is generally the only ESS and long-term equilibrium. Since defectors do not carry a cost, they always fare better in the absence of any measurable public good effect. Overall, the stability of the all-*D* equilibrium is independent of the functional form of the growth rates and solely depends on whether defectors have a higher overall intrinsic growth rate than cooperators. In contrast, the stability of the all-*C* state depends critically on the functional form of the intrinsic growth rates. For example, if *r*_*C*_ > *r*_*D*_ for all frequencies of cooperators in the population, then all-*C* is the ESS. A polymorphic ESS with a mix of *C* and *D* becomes possible when the ratio of respective growth rates is equal to ratio of death rates *r*_*C*_/*r*_*D*_ = *δ*_*C*_/*δ*_*D*_ for some frequency of cooperators. In these cases of coexistence, we are interested in the stable fraction of cooperators.

We label the fraction of cooperators in an equilibrium polymorphic state *y*\*, and the total population size in this state *Y*\*. When all baseline growth and death rates are the same for *C* and *D* (*α*_*C*_ = *α*_*D*_ and *δ*_*C*_ = *δ*_*D*_), the equilibrium population size is largest in the all-*C* state. A full linear stability analysis is presented in the Methods.

In the following, we assume that all baseline proliferation and death rates are equal, *α_C,D_* = *α* and *δ_C,D_* = *δ*. Then all possible differences in the intrinsic growth rates that could resolve the tragedy of the commons are determined by parameters in the frequency-dependent growth rates *r_C,D_*, such as the benefit parameters *β*, *σ*, and the cost term *κ*. In the case of a linear public good, *r*_*C*_ = *α*(1 + (1+ *β y* (*n* − 1))/*n*) − *κ* and *r*_*D*_ = *α*(1 + *β y* (*n* − 1)/*n*), there thus cannot be a polymorphic ESS. All-C is a stable ESS if and only if the benefit-to self of each cooperator exceeds the individual cost: *α β*/*n* > *κ*. Hence, smaller neighborhood sizes favor cooperation.

For a nonlinear form, e.g. sigmoidal growth rates, we still often observe just one internal equilibrium. In Figure 1, we give explicit examples of the possible phase diagrams that indicate whether a single polymorphic equilibrium exists and is stable. For fixed neighborhood size, baseline growth rates and cost of cooperation, these phase diagrams are uniquely determined by the two payoff parameters *β* and *σ*. Increasing the neighborhood size reduces the chances of observing a stable all-*C*. On the other hand, increasing the neighborhood size can increase the parameter range for which we can expect bi-stable evolutionary dynamics (unstable polymorphism).

Overall, a change in neighborhood size can critically alter the range of benefit parameters that allow for a stable equilibrium where cooperators and defectors coexist. For example a stable polymorphic equilibrium can be expected if the frequency-dependent benefit parameter *β* is an order of magnitude larger than the parameter *σ* that quantifies a frequency independent benefit, in which case the net growth rate advantage of cooperators is either non-monotonic or decreasing with increasing abundance of *C*.

### There are maximally two polymorphic equilibria, and only one is stable

How many polymorphic equilibria can we maximally observe? In the Methods we show that for the “almost identical” public good functions (*r*_*C*_ = *A r*_*D*_, for some constant *A* > 0) there can only be one internal equilibrium. The same holds for public good functions that are “always better” in terms of differential returns from the public good, e.g. a cooperator benefits more when another cooperator is added to the neighborhood, 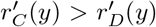 (see Methods). In general, a sigmoidal public good functions of the form in (3) can have a maximum of two polymorphic equilibria. One internal equilibrium is typically unstable and the other is stable. The stability properties of internal equilibria critically depend on the neighborhood size *n*. This means that saddle-node and transcritical bifurcations [36] are possible in which the neighborhood size *n* acts as the bifurcation parameter (Figure 2). The frequency-dependent benefit parameter *β* can also control such bifurcations.

An unstable and a stable polymorphic state can coalesce and annihilate each other at a critical point, known as a saddle-node bifurcation point. This event occurs without affecting the stability of the monomorphic states. Such critical points depend on specific parameter values of the public good function. Saddle-node bifurcations tend to co-occur with rather sharp transitions between no benefit (due to low numbers of cooperators) to a maximal benefit. Such sharp transitions can be observed for strong frequency-dependent selection (large values of *β*). In Fig. 2 A and B we show an example of *β* > *σ* leading to a non-monotonic growth rate differential, which creates a saddle-node bifurcation controlled by the neighborhood size. In contrast, for *β* < *σ*, we observe monotonically increasing growth rate differences, possibly leading to unstable coexistence and resulting in a transcritical bifurcation (Fig. 2 C,D) where the resulting ESSs are alternative stable states of all-*C* or all-*D*.

**Figure 1:**
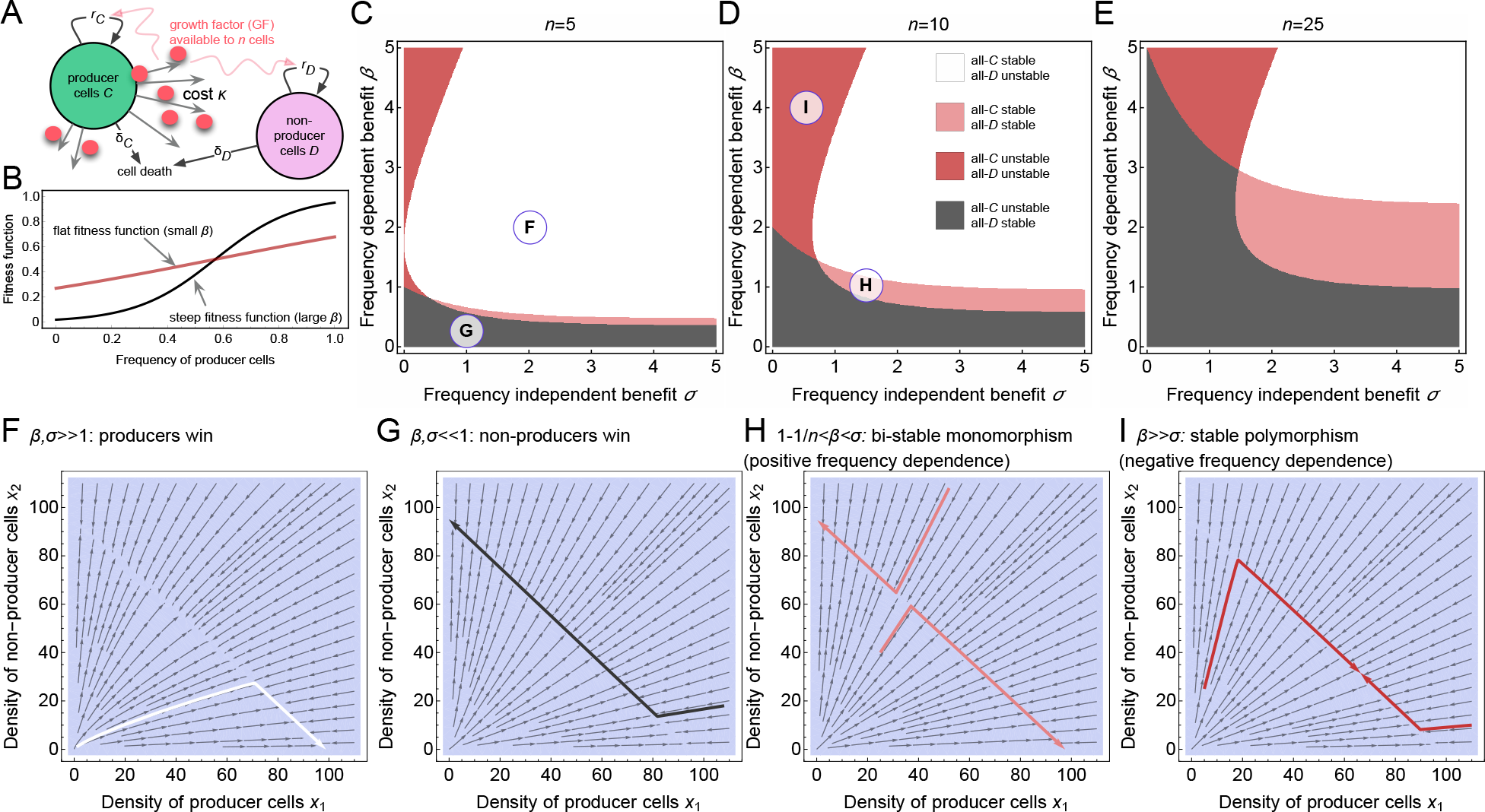
Benefit parameters and neighborhood size determine the direction of selection. **A**: Schematic of the Model, in which cooperators cells (type *C*) provide a public good-like growth factor (GF) at a cost, which benefits both cooperators and defectors (type *D*) within a defined neighborhood of *n* cells, resembling an effective diffusion range of the public good. **B**: Examples of nonlinear selection functions that determine the frequency-dependent selection effect on the intrinsic growth rates, (3b) with *n* = 8, *σ* = *β*/2, *β* = 2 (red) and *β* = 8 (black). **C-E**: Phase diagrams of the co-evolutionary dynamics for three different neighborhood sizes (given in the panels). Smaller neighborhood sizes favor producer cells. **F-I**: Stream plots (thin arrows) and example trajectories (thick arrows) of logistic co-evolution of *C* and *D* according to Eqs. (1), (3). The particular parameter settings of the previous panels are marked within their respective phase diagrams. In all panels we used *α*_*C*, *D*_ = 1.0/day, *δ*_*C*,*D*_ = 0.05/day, *κ* = 0.1/day, and *K* = 100, for different frequency independent, freq. dependent benefit and neighborhood size parameters; *σ* = 2.0, *β* = 2.0, *n* = 5, leading to dominance of the producers (**F**); *σ* = 1.0, *β* = 0.25, *n* = 5 (**G**), leading to dominance of the Consumers; *σ* = 1.5, *β* = 1.0, *n* =10 (**H**), leading to unstable polymorphism (bi-stability); *σ* = 0.5, *β* = 4.0, *n* =10 (**I**), leading to stable polymorphism (coexistence).

### Integrating empirical evidence of nonlinear cellular public good games

Public good functions that influence population growth rates in a nonlinear fashion have been proposed in the context of bacterial, yeast and cancer cell monolayers [31], and were subsequently measured empirically in an *in vitro* context [13]. However, a concise numerical and statistical interpretation of this data has been missing, partially because the proposed game theoretic framework to explain the data might have been too far removed from the biology of these particular cellular growth patterns [14]. With our model, we can in part close this gap and recapitulate the intriguing frequency-dependent cellular growth patterns measured by Archetti *et al.* (2015) [13] in cancer cells. They established that IGF-II (insullin-like growth factor 2) can act as a nonlinear public good that comes at a specific cost to cooperators (IGF-II producer cells). In the context of freely available nutrients (Fetal Bovine Serum; FBS), the benefit of producing the public good declines with increasing concentration of FBS (Fig. 2 E, see also Fig. 3C in [13]).

What are possible payoff functions in nonlinear cellular public goods games? We used data from cellular *in vitro* competition experiments using the population growth and expansion of IGF-II producer cells (cooperators) and non-producer cells (defectors). The actual neighborhood size was not measured or estimated for these experiments. Therefore, in fitting our model to this data, we considered neighborhood size to be independent and identically distributed between *n* = 4 (nearest neighbors in 2D) and *n* = 40 cells. For each value of *n* and for each experimental setting (percent FBS in the medium), we fit our mathematical modeling framework (Methods) to the cellular growth data, to estimate values for *β* and *σ*. Our estimated values of *β* ranged from 1.5 to 6 (Fig. 2 F), and values of *σ* from 0 to 3 (Fig. 2 G). For example, in the context of 5% FBS in the background medium, we observed median values of *β* = 3.67 and *σ* = 1.87. In our model, these parameter values would predict an unstable polymorphic equilibrium near *y*\* = 0.235, and a stable polymorphic equilibrium near *y*\* = 0.784 (Fig. 2 H). These estimates depend on the distribution of values of *n*, which highlights the need in future studies to devote more effort into determining the distance over which a public good spreads. Our statistical analysis shows that our model framework is capable of explaining the evolutionary dynamics of cellular games with nonlinear public goods.

### Type-dependent neighborhood sizes favors coexistence

Cooperators may experience a different neighborhood size, *n*_*C*_, than defectors, *n*_*D*_. Cooperators should be under selection to favor a smaller neighborhood size, and vice versa for defectors who should favor free-loading from as many individuals as possible. Thus, *n*_*C*_ ≤ *n*_*D*_. Here we examine how strategy-specific neighborhood sizes influence coexistence, the ESSs, and the eco-evolutionary dynamics (1). For this analysis, we assumed that cooperators interact with other individuals within their specific neighborhood of size *n*_*C*_, and that defectors interact within *n*_*D*_. Under these additional assumptions, we examine under what conditions a small population of cooperators can invade a resident population of defectors.

The relation *n*_*C*_ < *n*_*D*_, favors coexistence, which can be seen in the different bifurcation patterns in Figure 3. This is because each strategy experiences the neighborhood size most conducive to its success in a world composed primarily of the other strategy. When both *C* and *D* experience the same neighborhood size, then small sizes favor all-*C* and larger neighborhood sizes favor all-*D*. For a sufficiently small *n*_*C*_, *C* can invade a population of all-*D*. For a sufficiently high *n*_*D*_, *D* can invade a population of all-*C*. Hence *n*_*C*_ ≪ *n*_*D*_ reduces the likelihood that all-*D* and/or all-*C* will be an ESS, and increases the likelihood of coexistence via mutual invisibility. A mix of *C* and *D* can become the sole ESS.

### Fluctuations in neighborhood size

How do fluctuations in the size of the neighborhood of interacting individuals influence coexistence of cooperators and defectors? And how do feedbacks between total population size and neighborhood size influence the ESS of *C* and *D*? To analyze these questions, we devised a compartmentalized version of the eco-evolutionary growth model (1). Instead of exogenously fixing neighborhood sizes, we let the total population size influence neighborhood sizes by fixing the number of neighborhoods into *N* compartments. The total population of *C* and *D* individuals, *Y*, is then randomly distributed across these compartments, resulting in a multinomial distribution of neighborhood sizes with a mean of *Y*/*N*. In this model, the number of compartments acts as a surrogate for the inverse of an effective neighborhood size. This stochastic mixing of the population was performed numerically according to a multinomial sampling process (see Methods). Hence the interaction neighborhood for each strategy type was a stochastic variable determined by the number of compartments and the total population size. For example, for *Y* = 1000 and *N* = 100, the expected neighborhood size would be *Y*/*N* = 10. A schematic of the compartment-based dynamics is given in Fig. 4A, in which the neighborhood size is an emergent quantity. Of critical importance to the dynamics of this model is the time between the mixing steps (Fig. S1, S2, and S3). The details of this *N*-compartment modeling approach are given in the Methods section.

**Figure 2:**
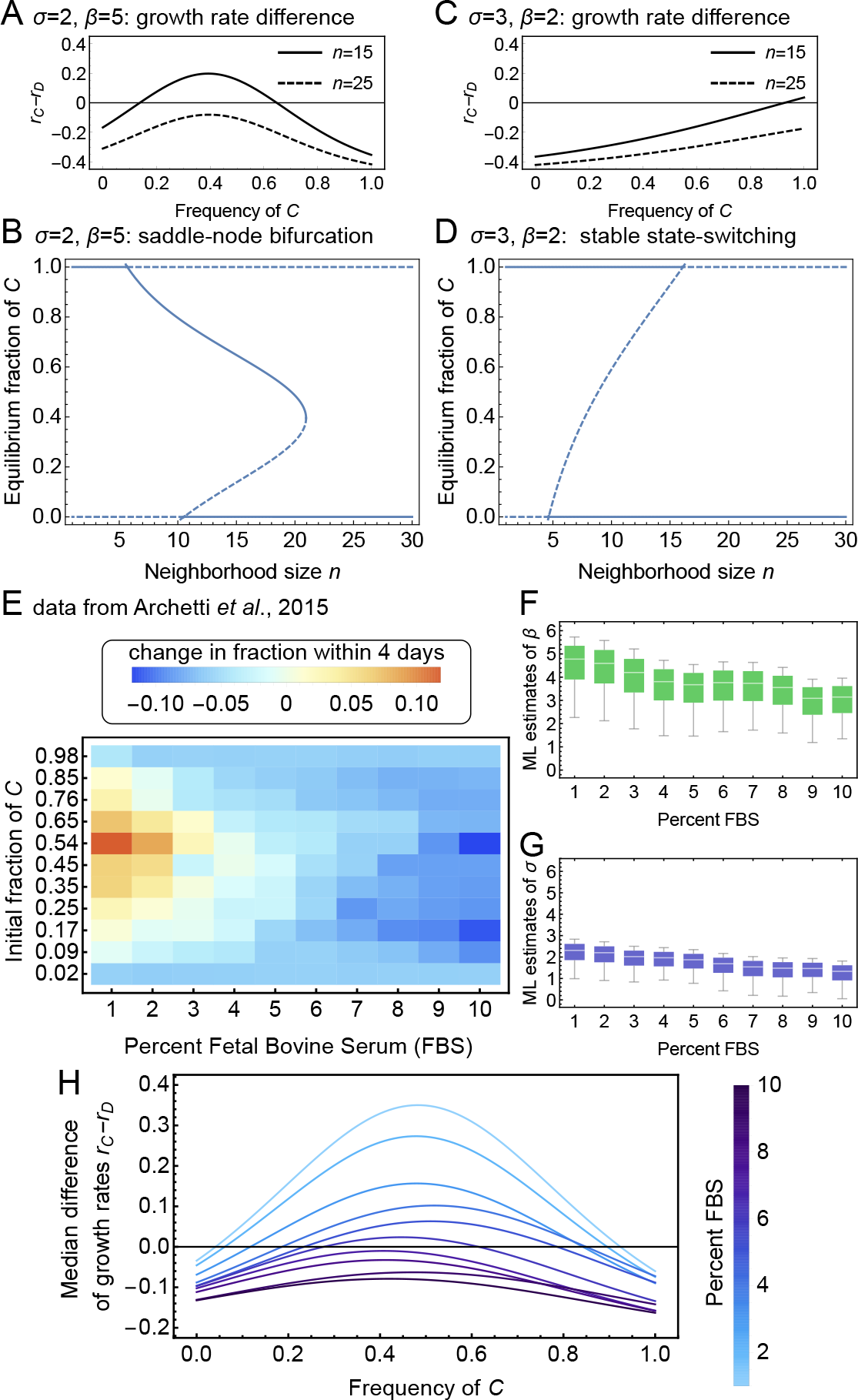
The nonlinear public goods function influences growth rate differences and invasion dynamics. **A** and **B**: For *β* > *σ*, differences in intrinsic growth rates can have a maximum at a fraction of cooperators between 0 and 1. The number and positions of equilibria then critically depends on neighborhood size, and saddle-node bifurcations are possible. **C** and **D**: For *β* < *σ*, differences in intrinsic growth rates tend to be monotonically increasing, and we can observe unstable coexistence and a form of a transcritical bifurcation pattern. For these model predictions we used *α_C,D_* = 1.0, *δ_C,D_* = 0.1, *κ* = 0.5, and a carrying capacity of *K* = 1000. **E**: Data on the frequency change of *C*, measured over four days by Archetti *et al.* (2015) [13]; using *β*-tumor cell lines derived from the Rip1Tag mouse model [37] with and without deletion of a gene responsible for production of the growth factor IGF-II, which was identified as the nonlinear public good. The two cell lines were grown *in vitro* with different concentrations of Fetal Bovine Serum (FBS). **F** and **G**: As neighborhood size, i.e. the group of cells playing the public goods game, have not been determined, we varied *n* from 4 to 40 cells and fitted a nonlinear model in form of the numerical solution of Eq. (5a), to obtain maximum likelihood (ML) estimates of *β* and *σ* for each FBS concentration. **H**: Growth rate differences of our re-analysis of Archetti *et al.*’s data (with a background growth rate set to 1/day, and using their estimated cost of public good production of *κ* = 0.25*α*, see Fig. 1 in [13], and an average neighborhood size *n* = 22). Low concentrations of FBS led to both stable and unstable coexistence, which was lost for higher FBS concentrations.

Public goods are produced and shared at the compartment level. A compartment’s contribution to the overall population growth rate of *C* and *D* individuals is based on the total population size across all compartments as a density effect (competition), and the number of *C* and *D* individuals within the compartment as the frequency-dependent effect. Neighborhood sizes become linked to the population dynamics via *Y*/*N*. The number of compartments remains fixed and hence the average neighborhood size increases linearly with total population size.

Each time step alternates population and frequency dynamics with a new round of random distribution of individuals among compartments. We observed that cooperators survived above a threshold number of compartments *N*, and that cooperators would become the ESS above a critical value *N*_crit_. Depending on the values of *β* and *σ*, coexistence of cooperators and defectors was possible. We observed the equivalent of a transcritical bifurcation (switching of stable states), controlled by an increasing number of compartments (Fig. 4 B), as well as the equivalent of a saddle-node bifurcation (Fig. 4 C) with stable coexistence at intermediate number of compartments. We defined the critical number of compartments such that above *N*_crit_ all-*C* would be stable. Figure 4 D shows how *N*_crit_ depends on the nonlinear public goods benefit parameters. Notably, if we did not re-initialize the population when changing the number of compartments, we could observe a strong hysteresis pattern (Fig. 4 E) as predicted by the transcritical bifurcation (compare this with Fig. 2 D).

**Figure 3:**
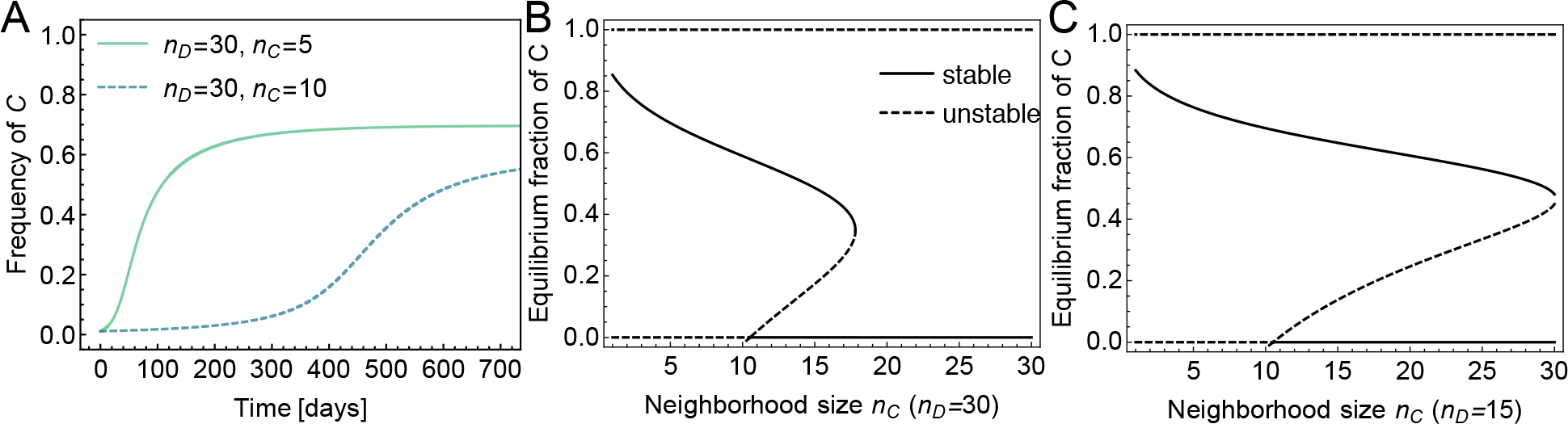
Differences in neighborhood size (public good sensitivity) can destabilize all-*C* but increase the coexistence regime. In all panels: *K* = 1000, δ_1_ = *δ*_2_ = 0.1, γ = 1.0/day, *σ* = 2.0, *β* = 5.0 and *κ* = 0.5/day. **A**: Temporal trajectories of the fraction of cooperators for two different values of *n*_*C*_, where we considered a resident population of defectors (*x*_*D*_ (0)/*K* = 1 − *δ/α*) that is invaded by a small population of cooperators (*x*_*C*_ (0) = 10), which thrive toward coexistence. The time to reach coexistence critically depends on *n*_*D*_ − *n*_*C*_. **B**: For fixed *n*_*D*_ = 30 the saddle-node bifurcation pattern of Fig. 2 B (where *n*_*D*_ = *n*_*C*_) changes such that the all-*C* state becomes entirely unstable, and we see a shift in the critical value of *n*_*C*_ above which cooperation is lost. **C**: Smaller values of, e.g. *n*_*D*_ = 15, act against defectors. Then we predict an increase in range of values of *n*_*C*_ for which cooperators and defectors stably coexist.

We can use a mean-field approach to analytically estimate *N*_crit_. Instead of using a multinomial distribution for the individuals among the *N* compartments, we let each compartment have the expected value, *Y*/*N*. We can then determine the number of compartments above which all-*D* becomes unstable and is no longer an ESS. Although this approach misses some important numerical and stochastic fluctuations, it recovers the general trend of the behavior of *N*_crit_ (Fig. 4 F), which shows non-monotonic behavior as a function of the public goods benefit. For small but increasing *β*, *N*_crit_ decreases monotonically, but then assumes a minimal value at intermediate selection strength, and then increases asymptotically toward the carrying capacity for very strong frequency-dependent selection (large *β*). These findings highlight the dynamical feedbacks between population dynamics influencing neighborhood sizes, neighborhood sizes influencing frequency dynamics, and frequency-dynamics influencing the population sizes. These feedbacks also point to how measurements of population densities and interaction distances could determine an effective neighborhood size.

## Summary and Conclusions

Eco-evolutionary dynamics include both changes in population size (population dynamics), and changes in the frequency of strategies within the population (frequency dynamics). Applying both to a public goods game we see how the two interact to impact the ESS. There is a critical role for neighborhood size. Neighborhood size describes the group of individuals that are close enough together to benefit from the public good. Our key results demonstrate the following facts: 1) smaller neighborhood sizes favors cooperators and increases the likelihood that all-*C* is an ESS, 2) a linear relationship between the frequency of *C* in the neighborhood and the overall value of the public good permits just two global ESSs, which are either globally all-*D* or globally all-*C* without the possibility of coexistence or alternate stable states, 3) a non-linear, sigmoidal relationship between the public good and the frequency of *C* allows for multiple stable ESSs, 4) under alternate stable states we observe saddle-node and transcritical bifurcations and hysteresis, and 5) feedbacks of population size on neighborhood size and/or strategy-dependent neighborhood sizes increases the likelihood of coexistence between *C* and *D*.

In a uniform environment, neighborhood size increases with the distance over which individuals experience the public good and with the packing density of individuals in the population. Social interactions may define the neighborhood as the members of a herd of antelope, troop of primates, or a web of colonial spiders. Here aggregation effects [38] may direct the production and distribution of the public good. Finally, neighborhoods may be formed when the distribution of suitable habitat exists as small patches within an otherwise inhospitable landscape. Widely spaced coral heads create smaller subpopulations of strongly interacting individuals (e.g. saddleback wrasse) that act as a neighborhood.

The inverse of neighborhood size is similar to the coefficient of relatedness in Hamilton’s rule [39]. Cooperation in a Prisoners Dilemma can be the ESS when relatedness is greater than cost divided by benefit. In the Prisoners Dilemma, the cooperator bestows the benefit on others while only incurring the cost. This lack of benefit-to-self is absent in the public goods game where the producer shares in the benefit. Despite this difference, the condition for cooperation (Prisoner Dilemma) and production (Public Goods Game) to be the ESS can be viewed as the same. In Hamilton’s Rule, the coefficient of relatedness represents the proportional return of ones actions via relatives, or neighbors. Analogously, in the public goods game, the inverse neighborhood (or group) size, 1/*n*, represents an individual’s share of its actions. When the benefits increase linearly with the frequency of producers, then cooperation is the ESS when 1/*n* is greater than the cost to benefit ratio.

**Figure 4:**
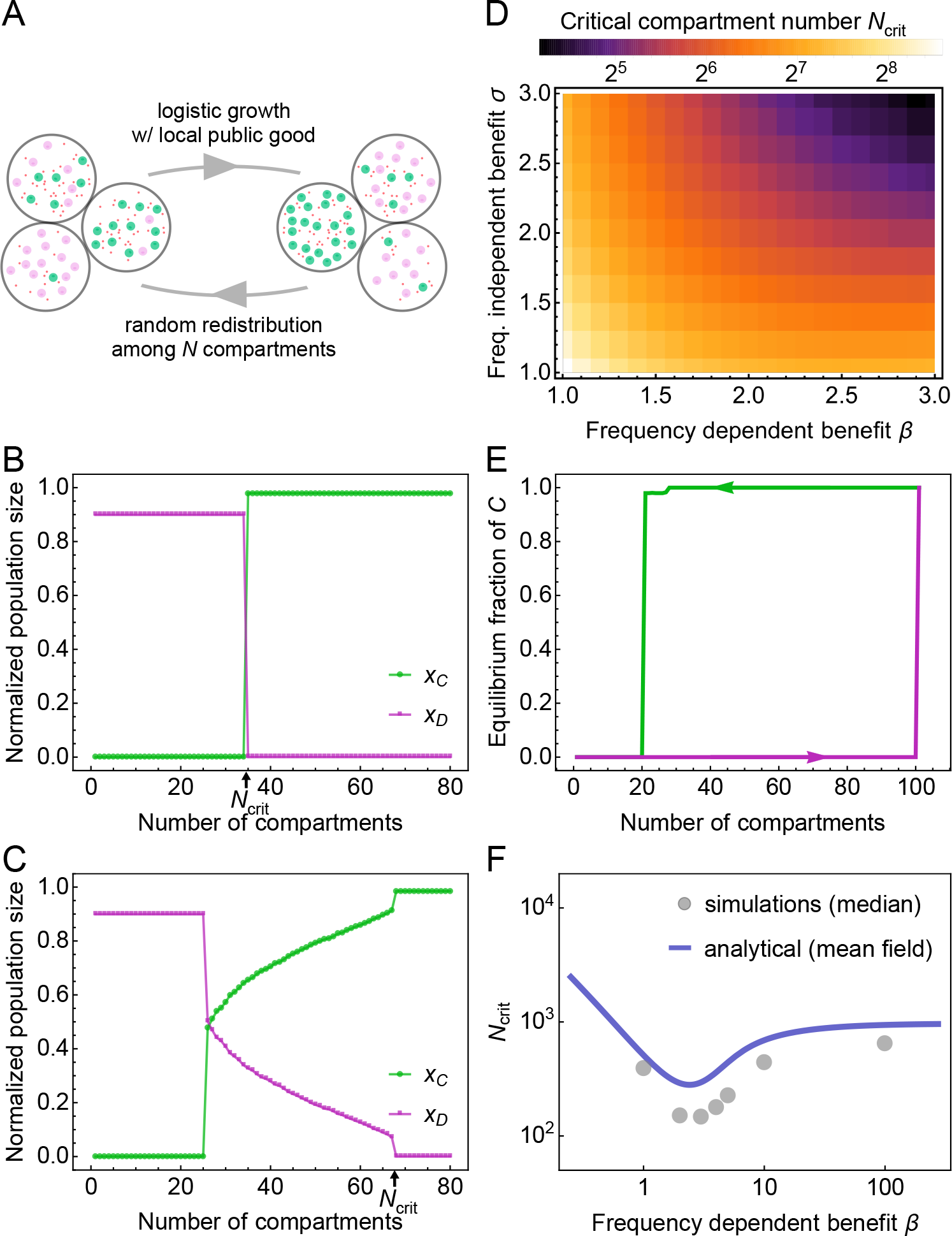
Fluctuating neighborhood size. can stabilize cooperation. **A**: Schematic overview of the quasi-spatial model. *N* compartments are filled with producers and consumers (a compartment can be empty). The system evolves from time *t*_0_ to *t*_0_ + Δ*t* according to the system (26). We repeatedly “mixed” (multinomial sampling *Y*-times into the *N* compartments) the system followed by selection for a time *T*_*S*_ = 10/α. **B**: For *β* < *σ* the system suddenly transitions to the all-*C* state with increasing number of compartments (decreasing effective neighborhood size). Parameters: *α* = 1, *β* = 2, *σ* = 3, *κ* = 0.5, *δ*_*C*_ = 0.1, *δ*_*D*_ = 0.1, *K* = 1000, *C*(0) = *D*(0) = 50, *T*_*s*_ = 10. **C**: For *β* > *σ* the system transitions to coexistence before equilibrating to the all-*C* state with increasing number of compartments (decreasing effective neighborhood size). Parameters: *α* = 1, *β* = 5, *σ* = 2, *κ* = 0.5, *δ*_*C*_ = 0.1, *δ*_*D*_ = 0.1, *K* = 1000, *C*(0) = *D*(0) = 50, *T*_*s*_ = 10. **D**: Calculation of steady state and location of *N*_crit_ for small *β*. The system transitions directly to the producer-only state. Parameters: *α* = 1, *κ* = 0.5, *δ*_*C*_ = 0.1, *δ*_*D*_ = 0.1, *K* = 1000, *C*(0) = *D*(0) = 50, *T*_*s*_ = 10. **E**: Hysteresis could be observed when changing *N* without re-initializing the population (parameters are the same as in **B**). **F**: Predicted *N*_crit_ using mean-field theory compared to the numerically obtained points. Parameters: *α* = 1, *σ* = 1, *κ* = 0.5, *δ*_*C*_ = 0.1, *δ*_*D*_ = 0.1, *K* = 1000, *C*(0) = *D*(0) = 50, *T*_*s*_ = 10.

The number of individuals comprising a neighborhood may be influenced by the total size of the population [40]. In this way, population dynamics can feed back on neighborhood size. In the case of alarm calls and prairie dogs there may be a direct relationship, as individuals must be within ear shot of each other. If neighborhood size increases with population density, then density feedbacks can result in the coexistence of cooperators and defectors. As a concesquence, an all-*C* world promotes a higher population size than an all-*D*, as it also allows a higher carrying capacity [28]. With a feedback on neighborhood size, an all-*C* population promotes a higher neighborhood size which, in turn, favors the invasion of defectors. Similarly, an all-*D* population promotes smaller neighborhood sizes that facilitates the ability of cooperators to invade. Density feedbacks have been shown to influence the ESS in games of spite [41] and producer-scrounger games [42]. However, if the public good is an essential nutrient secreted by a protist or cancer cell, then it may only provide resources for a fixed number of others, regardless of distance from the producer-so there may be little feedback at all. In many cellular interactions, we expect that a fixed number of nearby neighbors become the beneficiaries of a producer.

Linear public good games do not allow for saddle-node bifurcations, and consequently all-*C* or all-*D* are globally stable ESSs, unless there are density-feedbacks as discussed above. We show that a non-linear relationship between the public good and the frequency of producers does permit bifurcations and hysteresis and population diverse equilibria. We considered a sigmoidal relationship between benefit and cooperator frequency, which can favor coexistence because increasing returns at low fractions of cooperators act in their favor, whereas diminishing returns toward high fractions of cooperators favors defectors. We have observed that there are often maximally two interior equilibria, one will always be unstable and the other representing the stable coexistence of cooperators and defectors. With a different nonlinear relationship, such as a Michaelis-Menten (Hill) function (Eq. (11), see Methods), there can be up to five interior equilibria.

Three parameters controlled the eco-evolutionary equilibria: neighborhood size, the per cooperator size of the public good, and the upper limit on the total public good. These last two parameters interact in that a small benefit and large upper limit creates a near linear relationship between public good and cooperator frequency. A large benefit and small upper limit creates a near step function. Changing any of these parameters while holding the other two fixed can result in hysteresis whereby the shift from one equilibria state to another occurs at a different parameter value depending upon whether the parameter is being changed from small to large or vice-versa. For instance, as the benefit of the public good increases the ESS may shift from globally all-*D*, to either all-*D* or all-*C* as alternate stable states, and finally to globally all-*C*. Hence, there will be some range of benefit size where the ESS depends upon the initial population sizes of cooperators and defectors.

Cancer cells may engage in public goods games in a manner that make the disease more devastating and harder to treat [10]. Cancer cells produce signals that promote vasculature, or signal normal cells to produce this signal (vascular endothelial growth factor, VEGF). Then, the recruitment of blood vessels provides a benefit to the producer but also to any cells within some neighborhood of the new vessels. In such games, however, it is unclear whether there is a significant cost to cooperation. It may be possible to create therapies that incentivize the game in a manner that shifts the ESS towards defectors, for instance by decreasing the benefit of the public good. This approach may have demonstrable value in bicarbonate therapy, which is a means of neutralizing the low pH created by glycolytic cancer cells [43]. Increasing neighborhood size provides an intriguing, yet hitherto unappreciated therapeutic opportunity. Perhaps by increasing the diffusion rate of molecules (including VEGF) within the tumor the provider would experience less of a self-benefit and defectors would perceive a higher benefit.

Whether it be cancer or other ecological public good games in nature our work here suggests new ways to manipulate the parameters of public goods. Such manipulation could then achieve control, e.g. of pests or tumors, or promote polymorphism and diversity, e.g. of endangered or valuable species or cooperative sub-populations.

## Methods

### Co-evolution of cooperators and defectors in a public goods game

To begin, we rescale the populations *x*_*i*_ → *Kx*_*i*_ and so Eq (1) becomes

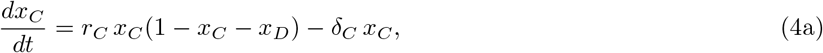

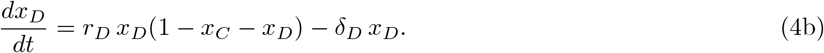

We map the dynamic system from producer/defector population to producer frequency and total population. We define *Y* = *x*_1_ + *x*_2_ and *y* = *x*_1_/*Y* and Eq. (4) becomes

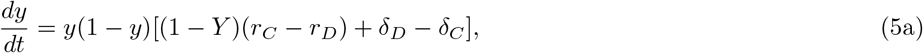

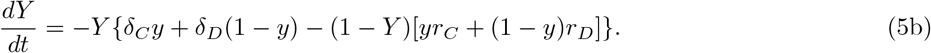

Based on the form given in Eq. (3), we analyze public good functions of the form *r*_*C*_ = *α_C_F_C_* − *κ* and *r*_*D*_ = *α_D_F_D_*, where we set *F*_*i*_(0) = 1, but we note a similar analysis can be done with *r*_*C*_ = *α*_*C*_ + *G*_*C*_ − *κ*, *r*_*D*_ = *α*_*D*_ + *G*_*D*_, where *G*_*i*_(0) = 0. The correspondence between the two forms is given by the relation *G*_*i*_ = *α*_*i*_(*F*_*i*_ − 1). We call *α*_*i*_ the public-good independent (intrinsic) growth rate.

### Equilibria points

The system governed by Eq. (5) admits a minimum of two fixed points: the boundary equilibria. The (*y*\*, *Y*\*) are given by

- *Defectors win*: 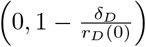.
- *Cooperators win*: 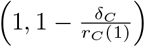.

Internal equilibria can exist provided that two conditions are met. First, both death rates are either both zero or both nonzero. If *δ*_*C*_ = *δ*_*D*_ =0, then coexistence is achieved with *Y*\* = 1 and *y* is undetermined (a line of non-isolated fixed points). If we suppose that *δ*_*C*_, *δ*_*D*_ are both positive, then we arrive at the *coexistence condition*

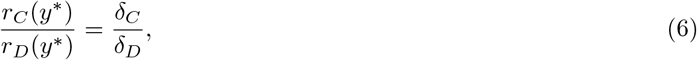

where 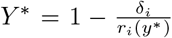. Finally we note that all physical trajectories (*y* ∈ [0,1] and *Y* ≥ 0) eventually enter the unit box [0,1] × [0,1]. To see this we only need to note that if *Y* > 1, then both terms in Eq. (5b) are positive and hence *Y* < 0.

### Linear stability analysis

#### Boundary equilibria

We analyzed the linear stability of the defector-only state first and obtained the Jacobian

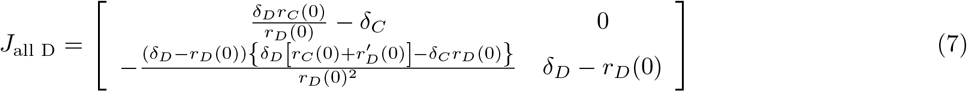

The eigenvalues of this system are λ_1_ = *δ*_*D*_ − *r*_*D*_(0), 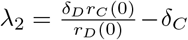. This state is stable provided both eigenvalues are negative, which leads to two conditions: (1) *r*_*D*_ (0) > *δ*_*D*_, and (2) *r*_*D*_ (0)/*δ*_*D*_ > *r*_*C*_ (0)/*δ*_*C*_. The first condition simply states the reasonable assertion that the intrinsic growth rate of defectors must exceed its respective death rate. The second condition states that the relative growth rate (the ratio of growth rate to death rate) of defectors exceeds that of producers.

The stability of the producer-only state is analyzed via the Jacobian

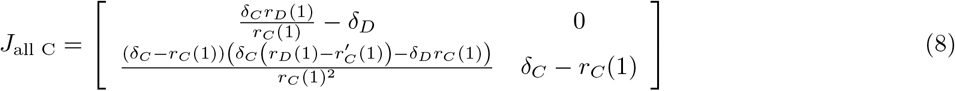

The two eigenvalues are λ_1_ = *δ*_*C*_ − *r*_*C*_(1), 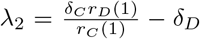. This state is stable provided that both eigenvalues are negative, which leads to two conditions: (1) *r*_*C*_(1) > *δ*_*C*_, and (2) *r*_*C*_(1)/*δ*_*C*_ > *r*_*D*_(1)/*δ*_*D*_. The first condition simply states the reasonable assertion that the intrinsic growth rate of producers must exceed its respective death rate. The second condition states that the relative growth rate of producers exceeds that of defectors. Note the similarity in stability criterion between the two states.

#### Internal equilibria

What if both states are unstable? To answer this, we consider the function 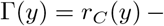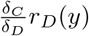. Then we see that the “all *C*” state is unstable provided that Γ(1) < 0 and the “all *D*” state is unstable provided that Γ(0) > 0. By the intermediate value theorem, there exists a *y*\* ∈ (0,1) such that Γ(*y*\*) = 0, which we recognize from Eq. (6) as the condition for a coexistence point. This shows the perhaps unsurprising fact that unstable boundary points in this system necessitate the existence of internal equilibria. It does not prove that the coexistence point is *stable*. Indeed, if there are stable closed orbits, a stable internal fixed point is not required. However, if the impossibility of closed orbits can be deduced, we can conclude that at least one internal fixed point must be stable. We will show later the impossibility of closed orbits in the case of equal death rates (e.g. *δ*_*C*_ = *δ*_*D*_ = *δ*).

It is useful to try to determine the maximum possible number of internal equilibria that can exist given a set of frequency-dependent growth rate function functions *r*_*C*_ (*y*), *r*_*D*_ (*y*). We are interested then in the number of possible zeros of Eq. (6), the *y* such that Γ(*y*) = 0. It is sometimes easier to locate extrema rather than zeros of a function. The correspondence between the number of zeros and extrema can be given by the following lemma:

#### Lemma

*Let P be the set of all coexistence points and E be the set of all extrema of the function* Γ. *The maximum number of coexistence points is given by* |*P*| = |*E*| + 1.

*Proof*. It is clear that a horizontal line can cross a function near an extrema at most two times. Ordering the extrema *E*_1_ < *E*_2_ <…< *E*_*n*_, we note that the interior extrema intersection points are counted twice. Hence, the number of possible coexistence points is given by |*P*| = 2|*E*| − (|*E*| − 1) = |*E*| + 1 which proves the claim.

With this lemma, the problem of determining the possible number of coexistence points is equivalent to determining the possible number of extremum of Γ. At this point we must specify a form of the public good function. We then investigate the sensitivity of the number of coexistence points to different public good functions. Extrema are located at the zeros of the function

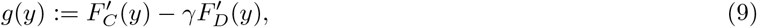

where we have defined 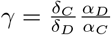. We analyzed the existence of internal equilibria for four types of functions with the following properties

- The public good always benefits the population (*F*_*i*_(0) > 1).
- More public good never hurts (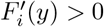 for all *y*).
- The good (unless linear) should eventually saturate (*F*_*i*_(*y*) → *F*_∞_ as *y* → ∞).

These three properties describe a many types of public goods. A notable exception was studied in the explanation of the Warburg effect, where the products of glycolysis are the public good used by cancer cells and at high quantities are actually harmful to the population [44]. First, consider the “almost identical” public good function *r*_*C*_/*r*_*D*_ = *A* = *const*. Eq. (9) becomes

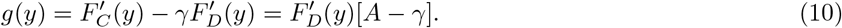

Since 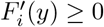, we conclude that *g*(*y*) is bounded away from 0 and hence there are no zeroes of *g*. This implies no extremum and so by the lemma there can be at most one internal equilibria. A similar result holds for the “always better” public good function. It is clear that if 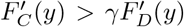 or 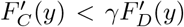, then *g* will have no zeros and we can conclude at most one internal equilibrium point is possible. These general results show immediately that the linear public good can never have more than one coexistence point and hence saddle-node bifurcations and other interesting phase diagrams are not possible. Note, these results are independent of the listed properties.

A general class of models which satisfies the three properties listed above are sigmoidal functions. We consider two general forms: the Fermi-like function considered in the main text, and the Hill function.

The Hill function

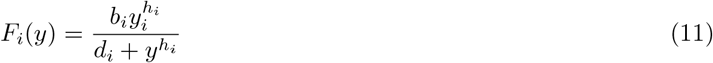

where the coefficients are all assumed to be nonnegative and *h*_*i*_ ∈ ℚ is the Hill coefficient. Plugging this into Eq. (9), and setting to 0 leads to

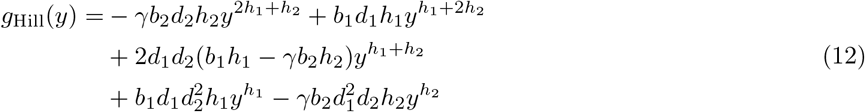

which we set to zero. There are three cases *h*_1_ > *h*_2_, *h*_1_ < *h*_2_ and *h*_1_ = *h*_2_. Let us investigate the case *h*_1_ > *h*_2_ first. Using Descartes’ rule of signs (DRoC), we obtain (−, +, sgn(*b*_1_ *h*_1_ − γ*b*_2_*h*_2_), +, −). Thus, DRoC shows that we can have at most four positive zeros if the middle term is positive and two if it is negative. The case *h*_1_ < *h*_2_ is analogous and leads to the same conclusion (with the condition on the middle term being reversed). The case *h*:= *h*_1_ = *h*_2_ is special since this combines many of the terms in Eq. (12): we require

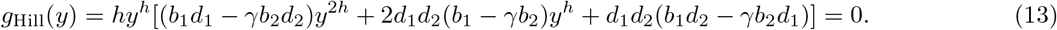

We can thus show that one cannot have alternating signs-hence we can only have at most one positive zero. Suppose the signs would alternate. Then we see that

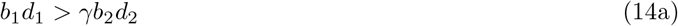

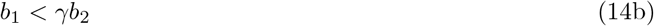

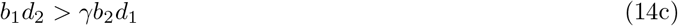

Eq. (14b) plugged into Eq. (14a) implies *d*_1_ > *d*_2_, while Eq. (14b) plugged into Eq. (14c) implies that *d*_2_ > *d*_1_, a contradiction. Hence, signs cannot alternate. To briefly summarize the results of the Hill function,

- If *h*_1_ ≠ *h*_2_, there are at most **five** internal equilibria.
- If *h*_1_ = *h*_2_, there are at most **two** internal equilibria.

Consider the Fermi benefit function 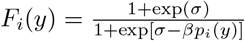 from Eq. (3) where *p*_*i*_(*y*) is the expected normalized benefit of the public good 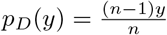 and 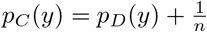. Plugging this into Eq. (9) gives

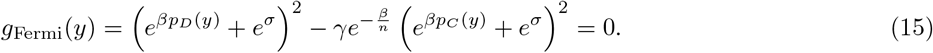

Simplification leads to

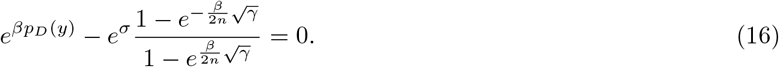

As before, this has a positive zero if the signs alternate. It is easy to show that the second term is always negative and so there exists a positive zero of this function. Hence, the Fermi function can have at most **two** internal equilibria. This should not be too surprising if we note the transformation of the Hill function via *y* = *e^x^*. The Hill function then transforms to

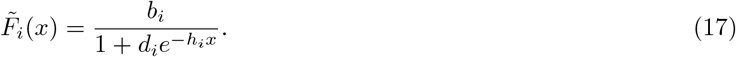

Letting *b*_*i*_ = 1 + *e*^σ^, *d*_*D*_ = *e*^σ^, 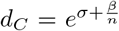 and 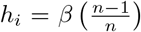 is the Fermi function used in the main text. Noting that a monotonic transformation conserves the location of extrema, we immediately see that the properties of the Fermi function actually follow as a corollary from the properties of the general Hill function considered. This is because *h*_*i*_ are equal and when *h*_1_ = *h*_2_ we showed that there can be at most two internal equilibria.

The general stability of the internal equilibria is much more complex. The Jacobian of an internal equilibria (*y*\*, *Y*\*) is given by

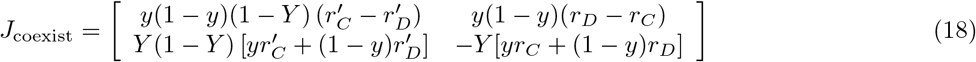

In this case, it is easier to look at the determinant Δ and trace τ. The values are

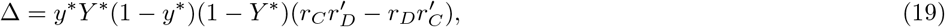

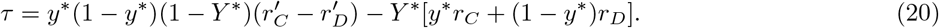

A necessary condition for linear stability of the internal equilibria is that Δ > 0, which is true if *r*_*C*_*r′_D_* > *r*_*D*_*r′_C_*, or using Eq. (6) and (9), *g*(*y*\*) < 0.

Let *δ* = *δ*_1_ = *δ*_2_. The coexistence points than occur at *r*_*C*_(*y*\*) = *r*_*D*_(*y*\*), with 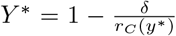. We note here that for physical solutions we require *Y*\* > 0 which implies that *r*_*C*_(*y*\*) > *δ*. The Jacobian is

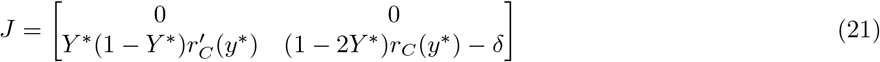

The eigenvalues are λ = 0, (1 − 2*Y*\*)*r*_*C*_(*y*) − *δ*. Based on the previous note of *Y*\* it is clear that the second eigenvalue is always negative and so all coexistence points are at least conditionally stable (there exists a trajectory which approaches the point). However, since one eigenvalue is 0, the nonlinear terms are relevant.

We want to find conditions that ensure stability of the coexistence points. Defining *u* = *y* − *y*\*, *v* = *Y* − *Y*\* and inserting these into Eq. (5) we obtain

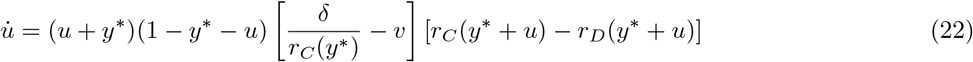

If we are close to the equilibrium (*u*, *v* ≪ 1), we can replace *v* = *Cu* where *C* is related to the entries in the Jacobian.

Plugging this into the above and expanding Γ_*i*_ in small *u* we obtain

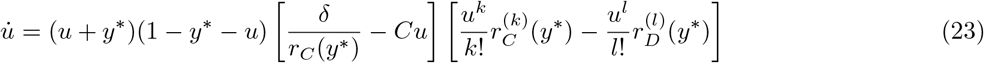

where 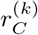,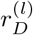 are the first nonzero terms in the expansion. Neglecting higher order terms we obtain

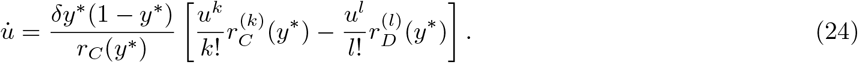

If *k* > *l* then the state is stable if 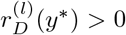, and if *k* < *l* then it is stable if 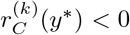. If *k* = *l* then the state is stable if 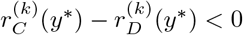.

### Impossibility of closed orbits

By index theory, we require at least one non-saddle fixed point to be in the interior of a closed orbit. Only interior equilibria can satisfy this requirement (the closed orbit cannot leave the unit box in phase space). Furthermore, if the closed orbit surrounds more than one interior equilibria, it must be an odd number 2*k* + 1 such that there are *k* +1 nodes and *k* saddles, with *k* ≥ 0. Consider first the special case when *δ*_*C*_ = *δ*_*D*_ = *δ*, then Eq. (5) reduces to

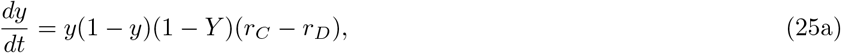

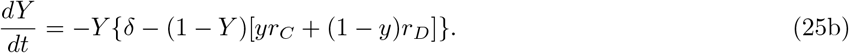

The coexistence point condition reduces to *r*_*C*_ (*y*\*) = *r*_*D*_ (*y*\*). It is clear that *ẏ* = 0 and *Ẏ* = − *Y*[*δ* − (1 − *Y*)*r*_*C*_(*y*\*)] on the line *y* = *y*\*. If *Y* < *Y*\*, then *Ẏ* > 0, while if *Y* > *Y*\*, *Y* < 0. It is impossible for a closed orbit to not transverse the curve Φ. However, the line is also invariant (a trajectory cannot leave once it is on it). Hence closed orbits are not possible, since no trajectory can cross through the line *y* = *y*\*.

### Nonlinear model fit

We used equation (5a) with the assumption *Y* ≪ *K* ((1 − *Y*) ⤒ 1in the rescaled dynamical system) to perform a nonlinear model fit in Wolfram Mathematica 11.2 (*NonlinearModelFit* and *ParametricNDSolve*) to obtain maximum likelihood estimates of *β* and σ. We set α = 1/day as this would only alter the overall time scale, and not the measured growth rate differences. We also set *κ* = 0.25/day, as estimated from Fig. 1 in [13], where under highest growth rate concentration the median difference between non-producer growth rate and producer growth rate amounted to 25% of the intrinsic growth rate. This fitting procedure was applied for every value of FBS concentration in the *in vitro* growth medium, for every integer value of *n* between 4 and 40, to result in ‘distributions’ of values of *β* and *σ* as reported in 2 F,G. Note that we did not statistically determine the neighborhood size *n*.

### Quasi-spatial model analysis

To simulate the effects of *N*-compartments subject to periodic mixing, we considered the 2*N*-coupled system of ODEs given by

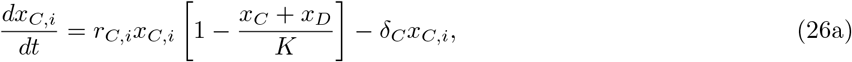

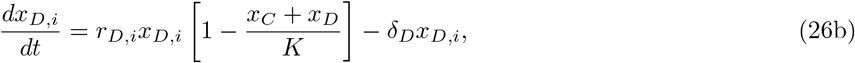

where *i* = 1,…, *N* and it is understood that 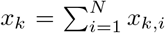. Each compartments dynamics are governed by the local interplay of producers and defectors within the compartment. The coupling between compartments is through the carrying capacity *K*.

Simulations were carried out in MATLAB according to the following scheme:

a. **Initialization**:

Set biological parameters (e.g. *α*, *β*, initial population size, number of compartments).
Distribute individuals among the compartments according to multinomial distribution.
b. **Dynamics loop**:

1. Integrate Eq. (26) from [*t*_0_, *t*_0_ + *T*_*s*_] for fixed *T*_*s*_.
2. If exit criterion is met, break out of the loop.
3. Redistribute cells according to the multinomial distribution. Go back to (1).
c. **Output**:

Dynamics of the total population (*t*,*x*_*C*_(*t*),*x*_*D*_(*t*)),
Final size (*x*_*C*_(*t*_final_),*x*_*D*_(*t*_final_)).

The exit criterion was met when *x_C_x_D_* = 0 for some *t* or if equilibration was reached. This was done by looking back a given amount of time steps and comparing the mean to the current value. After a set number of times they were within a certain tolerance, the loop would end. To solve the ODE, MATLAB’s ode45 built-in solver was used.

Aside from the compartments, all biological parameters are assumed fixed in the entire system (e.g. there is no compartment-dependent *β*). However, a new parameter − the selection time *T*_*s*_ is necessary. We investigated the impact of *T*_*s*_ for fixed compartment size. An interesting effect is observed in Fig. S S1. For *T*_*s*_ ⤒ ∞, mixing never takes place and in this scenario, the defectors go extinct. For finite *T*_*s*_, one observes coexistence. However, as T_s_ ⤒ 0 (time scales of mixing and selection are the same), the population appears to be moving towards fixation.

### Critical compartment size

It is clear that *N* is inversely related to the neighborhood size. Therefore, it is not surprising that there should exist some critical number of compartments at which the producers should be favored. To see this, note that *N* =1 is the original case where the neighborhood is the whole population. The defectors are the winners as this is the classic tragedy of the commons problem. On the other side, if *N* ⤒ ∞, each compartment will have at most one individual and hence the neighborhood size is effectively 0. In this case, the producers should win. We then expect there to exist some *N*_crit_ such that for *N* > *N*_crit_, producers are favored.

To approximate this, we use a mean-field approach where we replace the population size in each compartment by its expected value. We analyze the stability of the all producer population *y*_*i*_ = 1 for all *i*. We require the population size to be 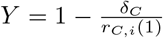. We replace *Y*_*i*_ by its expected value of *Y*/*N*. This collapses the *N*-compartment model to the original model and we recall the eigenvalues of the “all C” state λ_1_ = *δ*_*C*_ − *r*_*C*_ (1) and 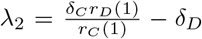.

Assuming the birth rate exceeds the death rate, the first is satisfied. The critical curve is then determined by λ_2_ =0. Solving this for the critical neighborhood size *n*_crit_, we obtain

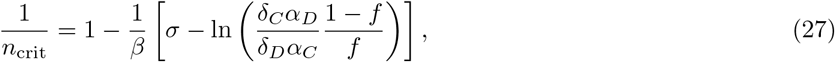

with

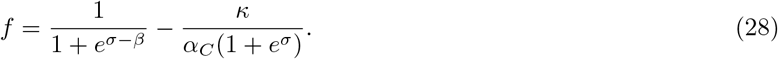

Now, the neighborhood size is the population size in the compartment and so its average will be 〈*n*〉 = *Y*/*N*. Plugging this into Eq. (27) yields (after scaling back carrying capacity)

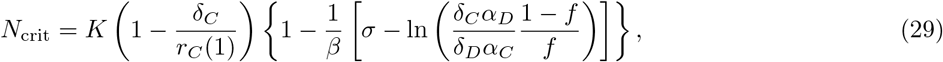

with

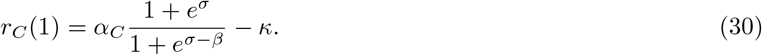

## Acknowledgements

We thank Marco Archetti (University of East Anglia, UK) for data sharing.

## Author contributions

P.G., J.S.B. and P.M.A conceived the project. G.J.K performed mathematical modeling. G.J.K and P.M.A. performed computational modeling. All authors analyzed the data and results, and wrote the manuscript.

## Competing interests

The authors declare no competing financial interests.

---

**Figure S1:**
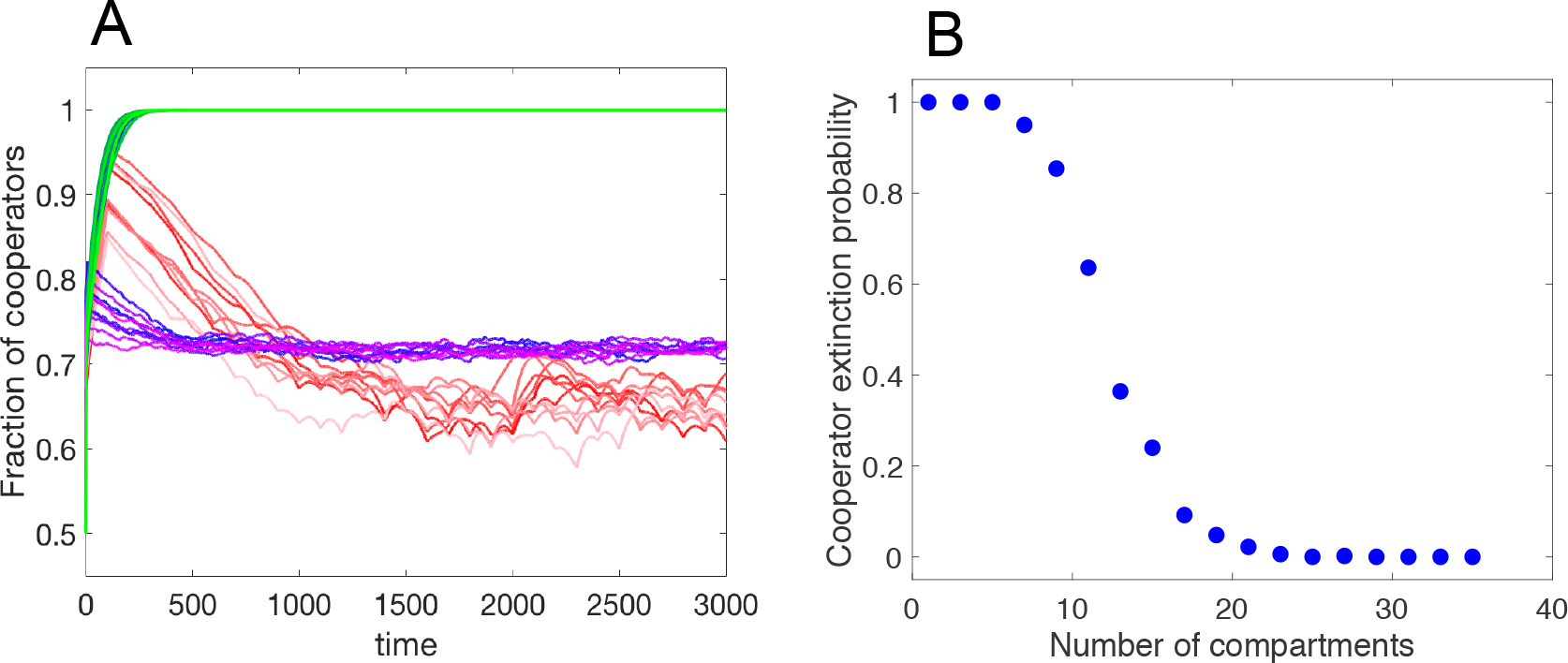
Stochastic effects of the *N*-compartment model. **A** The impact of selection time *T*_*s*_ = 10,100,1000, blue, red and green, respectively, with ten trajectories given for each. Parameters used *N* = 40, α = 1, *σ* = 2, *β* = 5, *κ* = 0.5, *δ*_*C*_ = *δ*_*D*_ = 0.1, *C*_0_ = *D*_0_ = 50. **B** *T*_*s*_ = ∞ (no redistribution after initialization). The producer extinction probability by number of compartments. Parameters used α = 1, *σ* = 3, *β* = 2, *κ* = 0.5, *δ*_*C*_ = *δ*_*D*_ = 0.1, *C*_0_ = *D*_0_ = 50.

**Figure S2:**
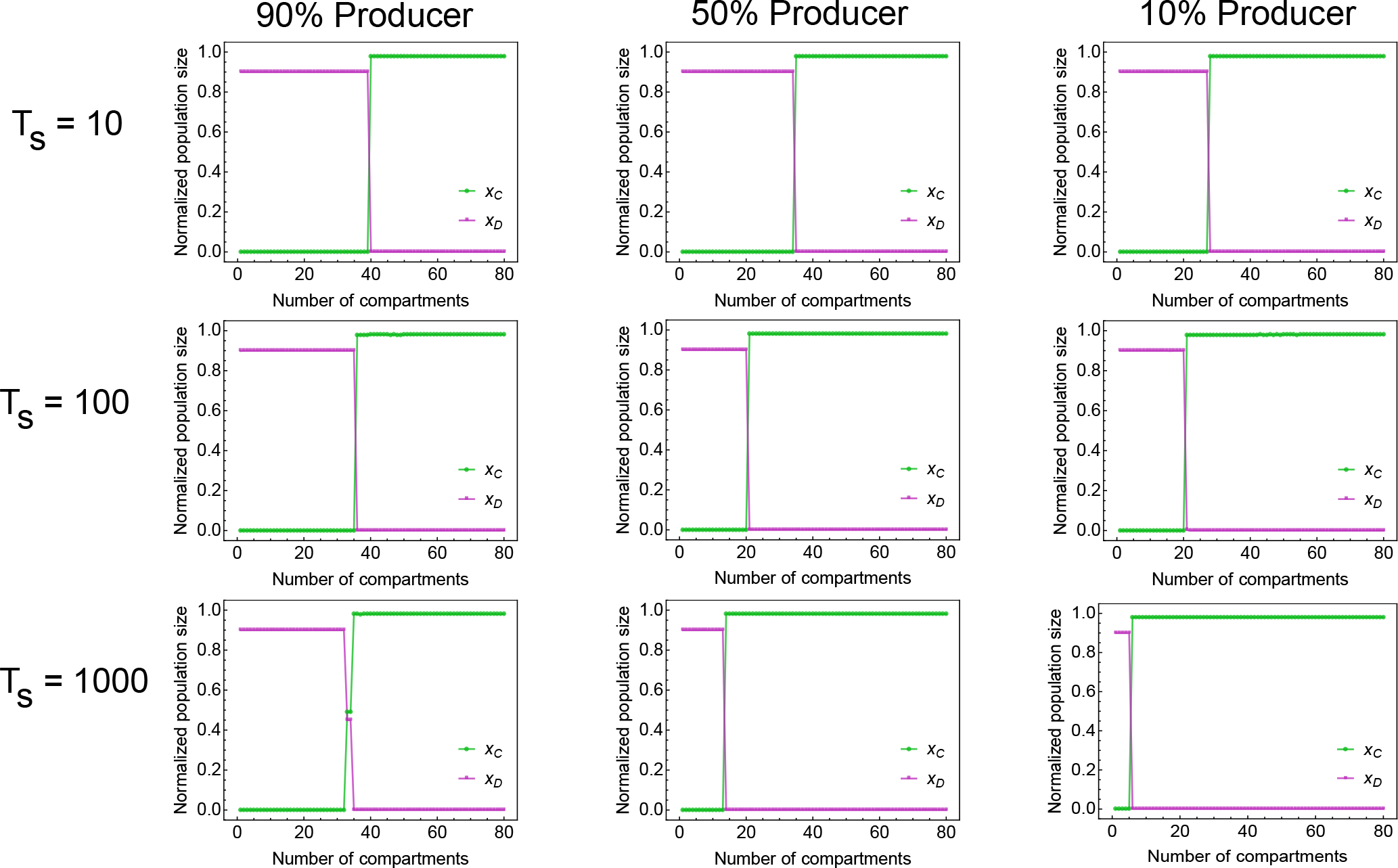
Critical compartment number and stable state-switching in the *N*-compartment model. Parameters: *β* = 2, σ = 3, α = 1/day, σ = 0.1/day, *κ* = 0.5/day, *K* = 1000.

**Figure S3:**
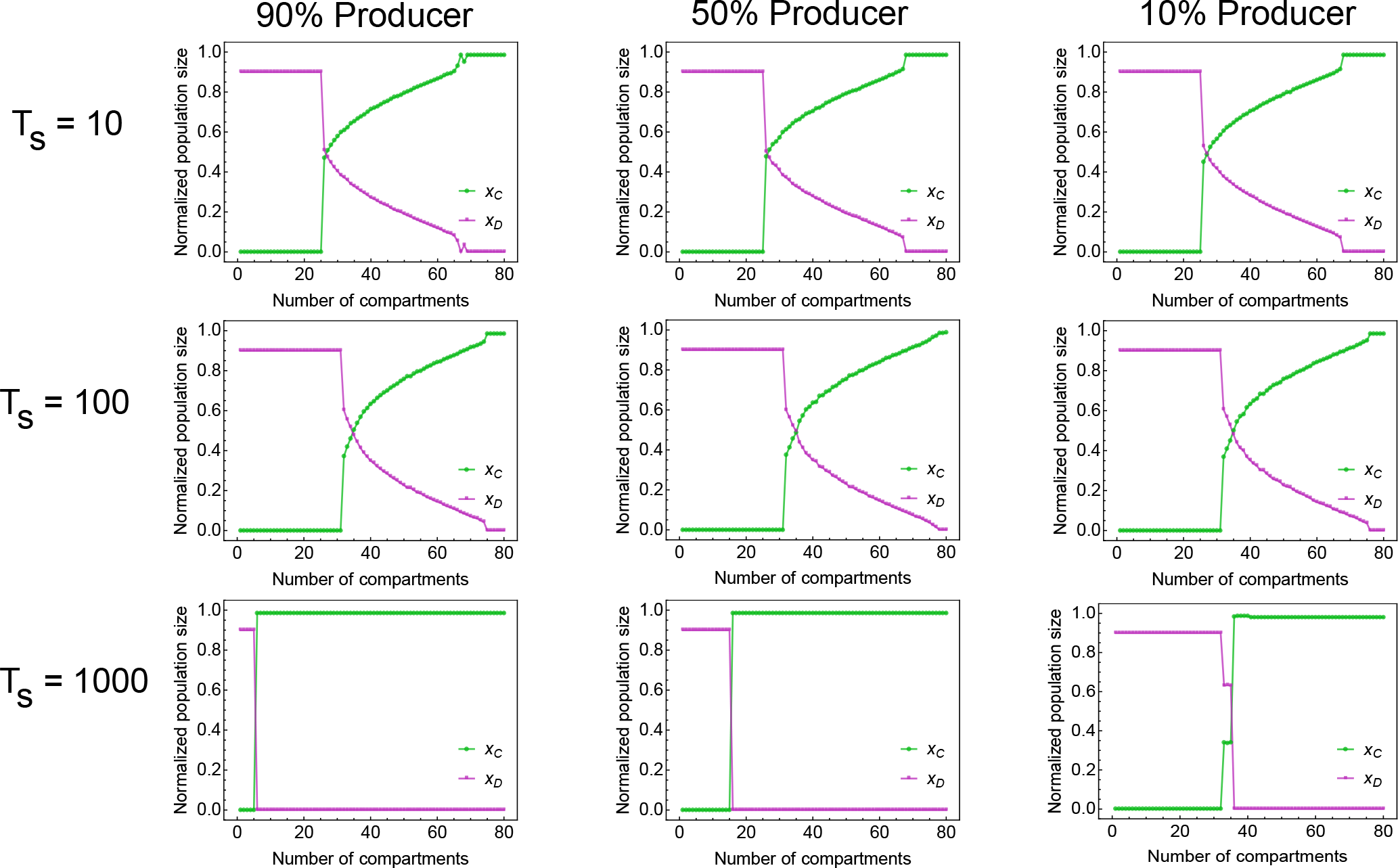
Critical compartment number and coexistence in the *N*-compartment model. Parameters: *β* = 2, σ = 3, α = 1/day, σ = 0.1/day, *k* = 0.5/day, *K* = 1000.

